# Global brain asymmetry and its variations in aging and related diseases

**DOI:** 10.1101/2024.08.29.610247

**Authors:** Haoyu Hu, Yi Pu, Yilamujiang Abuduaini, Xichunwang Wang, Clyde Francks, Paul M. Thompson, Xiang-Zhen Kong

## Abstract

Functional lateralization is a cardinal feature of human brain, and deviations from typical lateralization are observed in various brain disorders. Although this phenomenon has been widely acknowledged in the field of human neuroscience, decades of research have shown that it is a challenge to bridge the gap between (a)typically lateralized functions and hemispheric differences in structure (termed structural asymmetry). To address this important question, the present study employed the state-of-the-art machine learning techniques to investigate the brain structural asymmetry and its associations with cognitive functions, aging, and aging-related diseases, by integrating large-scale datasets. Our proposed multivariate approach revealed previously unknown and substantial structural differences between the left and right hemispheres, and established the associations between the global brain asymmetry and lateralized functions including hand motor and emotion processing. Furthermore, at the population level we mapped the aging trajectories of the global brain asymmetry, and unveiled significant diagnosis-specific variations in patients with Alzheimer’s disease and Parkinson’s disease, and individuals carrying a relevant genetic risk for atypical brain aging (i.e., APOE4 carriers). These results demonstrated left-hemisphere-linked changes in aging, which has challenged the traditional “right hemi-aging” model, and offered a promising approach for assessing brain aging and related diseases. Overall, our study with a novel approach presents one of the largest-scale investigation of global brain asymmetry, and takes an important step forward in understanding the intricate interplay between structural asymmetry, lateralized functions, and brain aging in health and disease.

**Significance statement:** Functional lateralization is fundamental to the human brain, with deviations linked to various brain disorders. Bridging the gap between functional lateralization and structural asymmetry has been a long-standing challenge. Using advanced machine learning and large-scale datasets, this study introduced a multivariate global brain asymmetry approach and revealed previously unidentified structural differences between the brain hemispheres, correlating these with cognitive functions, aging, and diseases like Alzheimer’s and Parkinson’s. Contrary to the traditional “right hemi-aging” model, we found left-hemisphere-linked aging changes. This work provides new insights into brain asymmetry, lateralized functions, and aging, offering a promising approach for assessing brain health and disease.

**Classifications:** Biological Sciences (Psychological and Cognitive Sciences);

## Introduction

One of the most prominent and evolutionarily conserved features of brain organization is its division into left and right hemispheres^1–3^. This division provides the brain with a greater surface area which indicates more “processing power”^4^, and is regarded to provide the structural basis which allows each hemisphere to develop functional specializations (i.e., functional lateralization)^5,6^. Lateralization is considered to be required for optimal functioning of the brain, and altered lateralization has been found to be associated with numerous cognitive and neuropsychiatric disorders, including dyslexia^7^, autism^8^, obsessive compulsive disorder^9^, and mood disorders^10^ (see also ^11,12^). Although the critical role of lateralization has been widely acknowledged in the field of human neuroscience, decades of studies have shown that it is challenging to link functional lateralization with hemispheric differences in structure (i.e., structural asymmetry)^13^.

Specifically, although the left and right hemispheres showed significant differences in behaviours and brain functions^14^, yet only minimal inter-hemispheric differences have been observed in structure indexed by various metrics such as cortical thickness and surface area^1,15^. For instance, over 90% individuals are right handers (dominant in the left hemisphere)^16,17^ and nearly 90% individuals demonstrate left hemisphere dominance in language tasks^18^. Variations in structural and functional asymmetry are only weakly correlated or even uncorrelated^16,19–21^. The lack of structure-function correlation substantially complicates the investigation of the structural basis of functional lateralization^3,11^. This important gap significantly hampers our understanding of the quintessential and human-unique organization of the human brain and severely limits the effective utilization of the organizational principle in assessing and diagnosing brain diseases.

Existing literature typically focuses on a localized analysis of inter-hemispheric differences of homologous regions (either at the vertex-, voxel- or region-level). However, this localized approach has revealed only minimal structural differences between the two hemispheres, and overlooked more complex patterns of the hemispheric differences^22^. This problem also applies to the existing studies of investigating altered asymmetry in brain disorders^11^. Such a methodological limitation motivated us to propose a multivariate perspective to study the inter-hemispheric differences (which we term “global brain asymmetry”). We reasoned that the functional hemispheric differences should manifest in global patterns rather than merely the constituent local differences^3,16,22^. Therefore, by taking multiple regions into account, it would allow us to describe the unique patterns of the structure of the hemispheres, thereby better revealing the complex intrinsic inter-hemispheric differences. Prior research offers corroborating evidence for our perspective that hemispheric differences should manifest in global patterns. One relevant example is brain torque, an overall hemispheric twist describing protrusions of the right frontal and left occipital regions over the midline, also known as “petalia”^3^. Another example is from recent large-scale studies, which have revealed a similar global asymmetry pattern in cortical thickness along the anterior-posterior axis^16,23^. A recent work of gray matter volume revealed multiple dissociable brain structural asymmetry patterns which also included a global pattern of brain torque^22^. These results converge to the idea that it is promising to explore the structural basis of functional lateralization from the multivariate perspective, instead of from the traditional univariate (localized) perspective.

Moreover, to our best knowledge, it remains a challenge to translate insights of the hemispheric differences gained from fundamental research into practical application such as in assessing healthy brain aging^11,24^ and diagnosing related diseases like Alzheimer’s disease (AD)^25,26^ and Parkinson’s disease (PD)^25,26^. Although previous studies suggest that lateralized functions change with aging^27,28^, yet those studies were usually based on behavioural observations with small sample sizes which resulted in different models and inconsistent results. For instance, the “right hemi-aging” model proposed by Dolcos et al. (2002), mainly based on behavioural observations, proposes that the right hemisphere shows greater age-related decline than the left hemisphere^27^. However, Cabeza (2002) reviewed existing studies and proposed a model called HAROLD (Hemispheric Asymmetry Reduction in OLder aDults), which indicates that cortical activation in cognitive functions of the brain tends to be less lateralized with aging (i.e., reduced asymmetry in cortical activation)^28^. A recent study reported interesting aging patterns of local hemispheric differences in regional cortical complexity across adulthood^29^. Also, altered asymmetries has been linked to aging-related diseases^30,31^ and existing studies suggest that asymmetric atrophy is evident in AD^3,32,33^. However, these findings remain to be replicated in independent and large-scale samples. Nevertheless, they provide supporting evidence for the critical role of brain asymmetry in aging. As reasoned above, our proposed approach might be more likely to capture the differences between the hemispheres compared to traditional approaches used in these studies. We therefore proposed to exploit the global asymmetry approach to chart the aging trajectories in the brain asymmetry and further explore the disease-linked variants. The results, in return, could provide additional support for the functional significance of our proposed global brain asymmetry.

In the present work, we employed advanced machine learning techniques to model the left-right hemispheric differences of the cerebral cortex from a multivariate perspective and subsequently explored the functional relevance of the global brain asymmetry. Specifically, we first applied an ensemble machine learning framework^34^ in a young-adult dataset (Human Connectome Project [HCP]: N = 1113, 22-37 years old) and trained predictive models based on morphological metrics (i.e., cortical thickness, surface area, and volume) for classifying the two hemispheres. We also explored the functional relevance by integrating the modeling results and a functional annotation analysis with a large-scale neuroimaging meta-analytic database (see Materials and Methods). To further investigated the links between the global brain asymmetry and aging-related diseases (e.g., AD and PD) at the population level (see Results), we next applied the proposed models to a large-scale aging cohort (i.e., UK Biobank; N = 43,919, 44-82 years old). Both individuals with healthy aging and patients with AD or PD were included. The prediction accuracy at different ages reflects aging-related changes in the brain-wide asymmetry patterns, and variations of the predicition accuracy in patients indicate deviations of the multivaraite brain asymmetry from the young adults. Finally, we examined the effects of relevant genetic variants on the aging of the global brain asymmetry to explore the genetic correlates of the deviation observed in the aging-related diseases. Common variants in the apolipoprotein E (APOE) gene are well known to be associated with AD, and perhaps also with cogntive and motor decline in PD^35–38^. Thus, we investigated whether individuals who were not diagnosed with either AD or PD, but with different APOE genotypes, would show different aging-related changes. Collectively, our study established a link between structural asymmetry, lateralized functions, and brain aging in health and disease, as well as the potential genetic correlates underlying this link.

## Results

### Multivariate modeling of hemispheric differences

We proposed a new approach to investigate the inter-hemispheric differences from a multivariate perspective. In this approach, we integrated advanced ensemble learning techniques to model structural differences between the two hemispheres of the human brain.

Specifically, we first developed and validated our approach using the HCP dataset which consists of high-quality brain structural images from a large number of young adults (N = 1113; 22-37 years old) (see Materials and Methods). We applied PipelineProfiler to visualize the best ensemble model from several base models created by *auto-sklearn*^39^ (Fig. 1A-1B; Fig. S1, S2). Morphometric measures of 34 cortical regions of each hemisphere were input to the models as features for that hemisphere. Models were trained and validated for each of the morphometric measures (i.e., cortical thickness, surface area, and volume) separately, resulting in three prediction models. All three models showed that the left and right hemispheres could be distinguished with high classification performance, i.e., cortical thickness: accuracy = 87.75%, surface area: 100%, and volume: 99.02% (Fig. 1C-1E). Each of the models was an ensemble of multiple branches/submodels with different classifiers, feature preprocessors and balancing strategies (see Materials and Methods). Single submodel with the highest ensemble weight within each of the validated models also showed high accuracy (i.e., 81.4% for Adaboost with Feature Agglomeration, 98.9% for Extra Trees with Fast ICA and 98.9% for Passive Aggressive with Kitchen Sinks for the three metrics, respectively) (Fig. 1B, S1B, S2B). The high performance indicated that the two hemispheres are definitively distinguishable from a multivariate perspective. Previous studies suggest very subtle differences between homologous regions (e.g., the largest Cohen’s d of 0.6 for cortical thickness asymmetry, indicating a larger than 76.4% overlap between groups for the majority regions)^11^. With the present dataset, the inter-hemispheric differences of homologous regions showed similar strength (see Supplemental Dataset S1-S3).

**Fig. 1.**
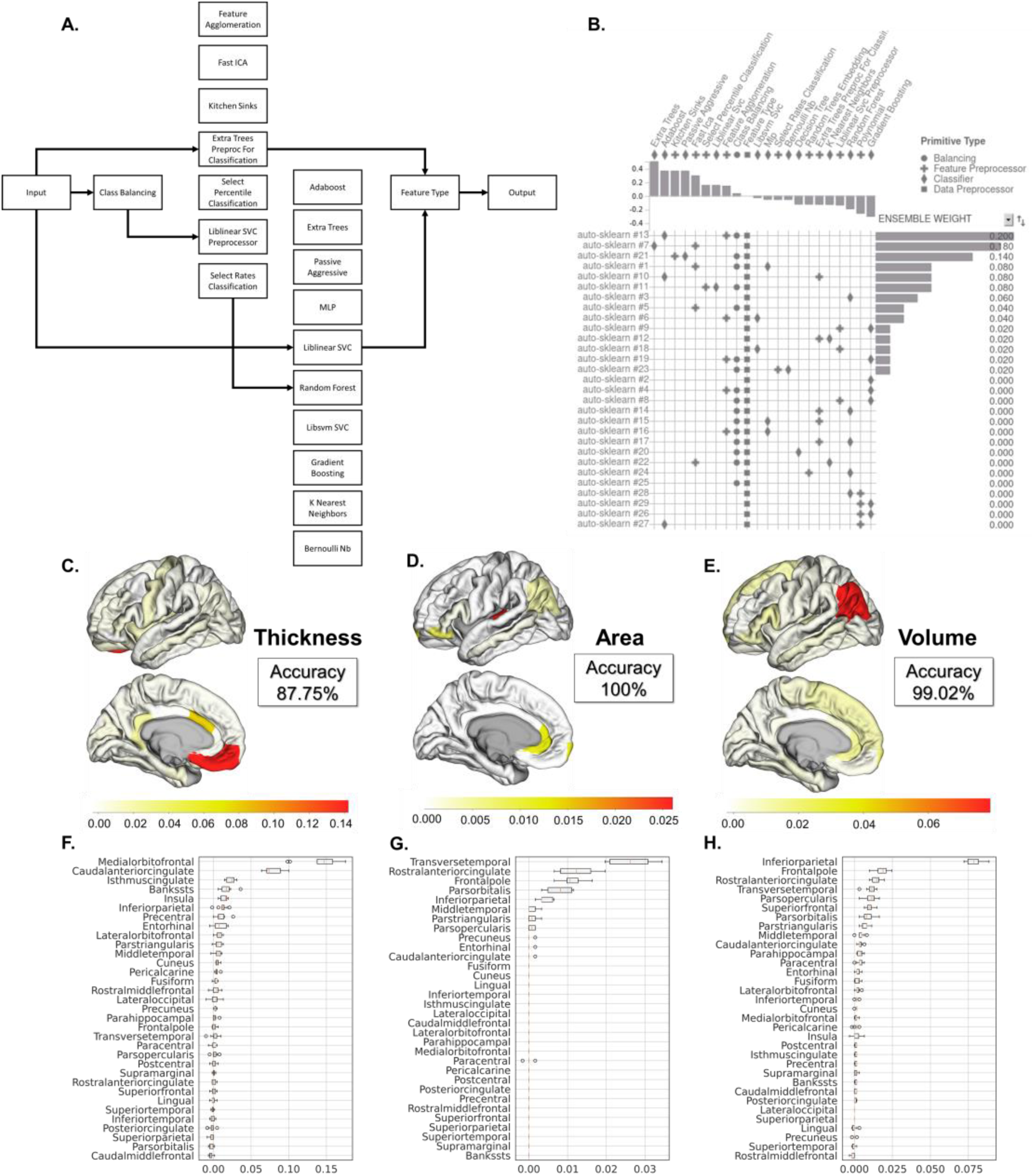
Multivariate modeling of hemispheric differences. (A) An example of the step-by-step autoML PipelineProfiler flowchart; (B) An example pipeline matrix (based on the cortical thickness model) with the top #1 classifier, which is called Adaboost. The matrix indicates primitives (data preprocessing, classifiers, feature preprocessor, and balancing applications; in the *columns*) used by each pipeline (i.e., each branch of the ensemble model; in the *rows*). Bar plots show the ensemble weight ranking (*left*) and the primitive contribution (correlation between primitive usage and performance contribution; *top*). (C) The importance scores of the cortical regions estimated in the cortical thickness model are illustrated in the brain space, and the box plot below indicates the importance ranking (F). (D) The importance scores estimated in the cortical surface area model are illustrated in the brain space, and the box plot below indicates the importance ranking (G). (E) The importance scores estimated in the volume model are illustrated in the brain space, and the box plot below indicates the importance ranking (H).

Based on the findings, we reasoned that the two hemispheres possess unique multivariate morphological patterns and that the substantial differences might have the potential to explain significant lateralization in cognitive functions and behaviours like handedness, language, and emotion processing. Therefore, we set out to test the functional relevance of the global brain asymmetry in a series of analyses below.

### Functional relevance of the global brain asymmetry

We explored the functional relevance by integrating the modeling results and a functional annotation analysis with a large-scale neuroimaging meta-analytic database (see Materials and Methods).

First, we estimated each region’s contribution in the models indexed by the importance score using a permutation-based approach (see Materials and Methods). It is important to note that the approach employed to estimate a single region’s importance in these models is distinct from approaches that rely on model coefficients, which can be considered questionable at times (e.g., ^40^). The results showed that different cortical regions showed various importance scores, and the importance score distribution across the cortex varied across the three models (Fig. 1C-1H). Specifically, regions such as the medial orbitofrontal region, the caudal anterior cingulate, and the temporal pole showed the highest importance scores in the model of cortical thickness; regions including the frontal pole, the *pars orbitalis*, the inferior parietal region, and the transverse temporal region contributed the most in the surface area model; regions including the frontal pole, the inferior parietal region, and the rostral anterior cingulate showed the highest importance scores in the volume model. Given the functional specificity of brain regions^41^, the different distributions of the importance scores across the brain may indicate that the three models have different functional relevance.

Next, we further identified the top functions related to each model using a functional annotation approach with Neurosynth, a well-established tool for interpretation of brain maps and having been widely used in the field (e.g., ^42–45^). Spearman’s rank correlation with a spatial spin permutation test (n=10,000) were employed for assessing the potential relationships and their statistical significance (see Materials and Methods). Results showed behavioural associations of the importance score maps with various functions (Fig. 2A). Specifically, the importance map of the cortical thickness model showed highest correlations with functional lateralization of motor-related functions (e.g., “motor task”: *r* = -0.52, *p* < 0.0001, FDR corrected); the surface area model showed highest correlations with functional lateralization of spatial attention and memory (e.g., “attention”: *r*= 0.51, *p* = 0.00005; “encoding retrieval”: *r* = 0.50, *p* = 0.00035); the volume model showed highest correlations with anxiety-, attention-, and memory-related functions (e.g., “anxiety”: *r* = 0.55, *p* = 0.0001; “attentional”: *r* = 0.44, *p* < 0.00035; “encoding”: *r* = 0.45, *p* < 0.0006). With group-level contrast maps of task fMRI from HCP (see Materials and Methods), we found that both the surface area model and the volume model showed significant correlations with motor and social processing (e.g., MOTOR_AVG-CUE: *r* = 0.48, *p* = 0.0007 for surface area, *r* = 0.35, *p* = 0.0018 for volume; SOCIAL_RANDOM: *r* = 0.35, *p* = 0.0045 for surface area, *r* = 0.58, *p* = 0.0002 for volume). Besides, the surface area model also showed a significant correlation with working memory (WM_AVG-TOOL: *r* = -0.36, *p* = 0.0051) and the volume model showed significant correlations with gambling (e.g., GAMBLING_ PUNISH: *r* = 0.48, *p* = 0.0014). In addition, the cortical thickness model showed a significant correlation with relational processing (e.g., RELATIONAL_MATCH-REL, *r* = 0.40, *p* = 0.0012).

**Fig. 2.**
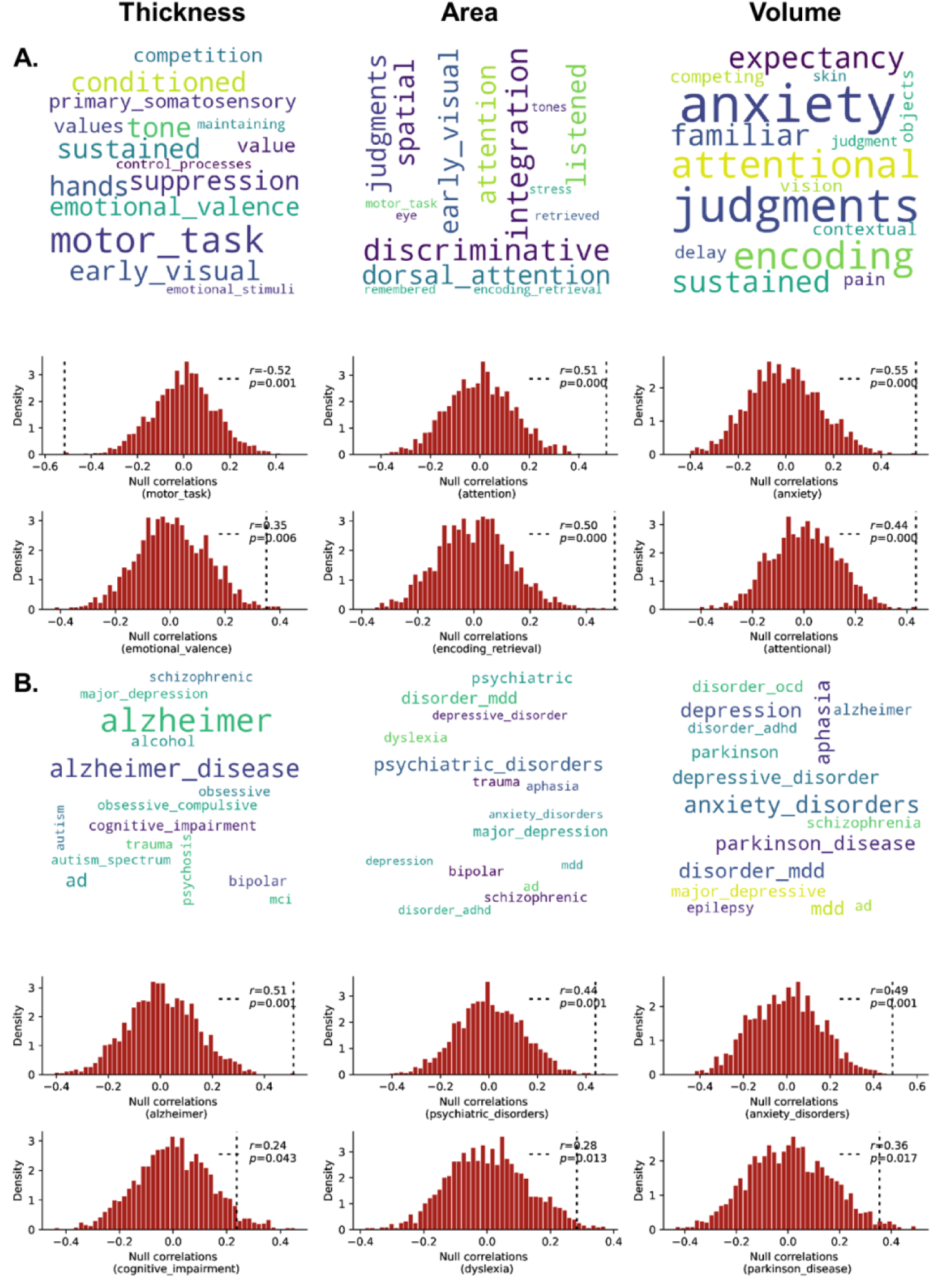
Functional annotation results of the prediction models of three morphological metrics (i.e., cortical thickness, surface area, and volume). Cognitive (A) and brain disorder (B) terms revealed in the functional annotation analyses are shown at the top. Font size indicates the strength of the correlation between the importance scores and the association maps relevant to each term of interest. Terms with the lowest 15 significance values are shown for visualization. Spin permutation test results of two exemplar terms are shown at the bottom, for each model.

Furthermore, we found various associations between the importance maps and lateralized atrophy in different brain disorders (FDR corrected) (Fig. 2B). Specifically, the cortical thickness model showed strongest correlations with AD (e.g., “alzheimer”: *r* = 0.51, *p* = 0.00025; “alzheimer disease”: *r* = 0.49, *p* = 0.0003), and the volume model showed strongest correlations with anxiety disorders and PD (e.g., “anxiety disorders”: *r* = 0.49, *p* = 0.0007), while the surface area model showed wide correlations with more general psychiatric disorders (e.g., “psychiatric disorders”: *r* = 0.44, *p* =0.0008).

Taken together, the series of analyses above provide supporting evidence for the functional relevance of the global brain asymmetry.

### Global brain asymmetry in aging and aging-related diseases

We noted from the analyses above that the global brain asymmetry showed significant associations with the lateralized atrophy in various aging-related diseases (e.g., AD and PD). To establish these links at the population level, we investigated the global brain asymmetry in aging individuals without any related disease and those with these aging-related diseases in a large-scale aging cohort (i.e., UK Biobank; 44-82 years old).

### Typical trajectories of the global brain asymmetry in healthy aging

First, we investigated how the global asymmetry changes during aging. Briefly, we trained a predictive model for each morphological metric for classifying the two hemispheres in a young adult cohort (i.e., HCP), and then applied the models to older individuals from the UK Biobank (see Materials and Methods). The prediction accuracy across different ages within the aging cohort may reflect aging-related changes in the global brain asymmetry patterns: for instance, a lower accuracy in increasingly older adults may indicate more deviation from the reference asymmetry pattern found in young adults.

Prediction results showed that the models for surface area (accuracy = 98.03%) and volume (83.59%) showed better overall performance compared to that of the cortical thickness model (66.88%) in the aging groups, which was in line with the performance in the training dataset. All the three accuracy scores were higher than the chance level (i.e., 50%). Interestingly, when summarizing the prediction results within each year of age, we found that the three morphological models showed different aging trajectories (Fig. 3A, 3F, 3K). Specifically, we found that the prediction performance with the cortical thickness and volume models showed a decreased trend with aging (cortical thickness: *r* = -0.58, *p* < 0.001; volume: *r* = -0.85, *p* < 0.001), suggesting continuous changes of the global hemispheric differences in aging. Results of curve fitting suggested quadratic and linear changes with aging for the cortical thickness model and the volume model, respectively (cortical thickness: quadratic model, *R*^2^=0.56, *F* = 19.12, *p* < 0.001; volume: linear model, *R*^2^ = 0.73, *F* = 93.39, *p* < 0.001) (Fig. 3A, 3K). Such a variability in prediction across different ages were mainly attributed to a rise in mislabeling of the left hemispheres (i.e., mistakenly labeled the left as the right) in the cortical thickness model (Fig. 3B). Similarly, an increase in mislabeling of the left hemisphere were found in the volume model (Fig. 3L). We term this phenomenon “left-hemi aging” (see Discussion). In the surface area model, we found that the predictive performance slightly increased with aging (cubic model, *R*^2^ = 0.57, *F* = 12.96, *p* < 0.001), particularly after 70 years old (Fig. 3F). Such effects were mainly attributed to less frequent mislabeling of the right hemisphere in the more elderly people (Fig. 3G). Simple correlation analyses of the asymmetry index across regions between the aging samples and the young adults showed similar but less pronounced effects (Fig. 3E, 3G, 3O) as the prediction results (Fig. 3A, 3F, 3K). We also calculated the inter-hemispheric differences of homologous regions within each age group, and the results showed various aging trajectories for different regions. A general decrease trend with aging was observed in cortical thickness while cortical surface area and volume differences of most regions were relatively stable (see Supplemental Dataset S1-S3 for more details).

**Fig 3.**
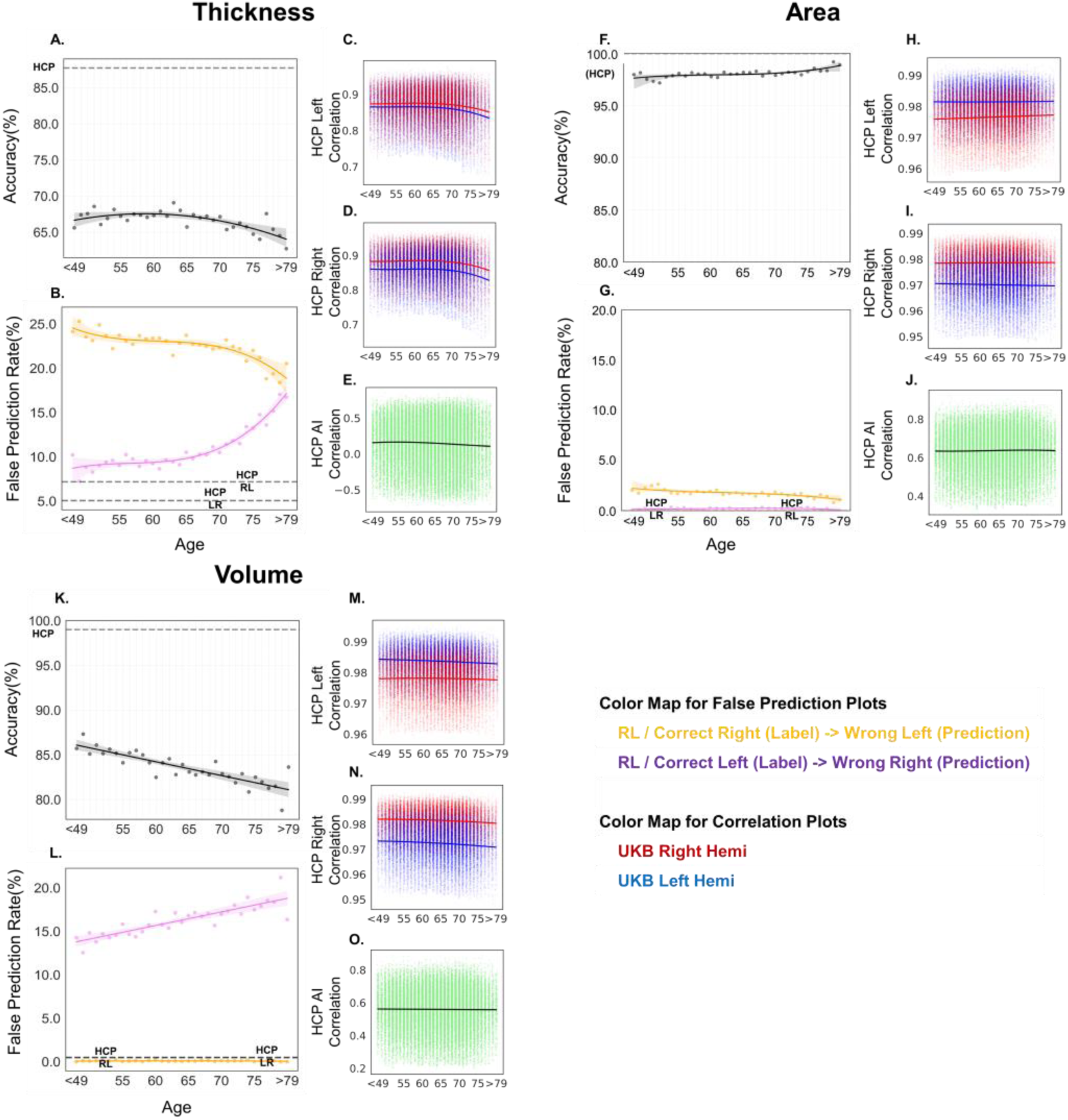
Multivariate asymmetry with brain aging. When applying the classification models trained in the young adult samples to the aging samples, the prediction accuracies vary at different ages: (A) cortical thickness; (F) surface area; (K) volume. (B), (G), and (L) show the false prediction rates of samples for each hemisphere. The orange color indicates the percentage of right hemisphere being mislabeled as the left, while the pure color indicates the rate of left hemisphere being mislabeled as the right. The side panels with scatter plots demonstrate the correlation-based similarity in the cortical measures of each hemisphere (left: C, H, M; right: D, I, N) and the asymmetry index (E, J, O) between the UKB samples and the HCP samples. These panels show how the global pattern in different cortical measures change with aging, taking the global patterns of young adults from HCP as the reference. Note that, considering the limited sample size of each age year of below 49 years old in the dataset, samples with an age of younger than 49 years old were grouped together when calculating the statistics (e.g., prediction accuracy). Similarly, samples of above 79 years old were also grouped.

As a sensitivity analysis, we additionally examined whether these observed aging-related changes in the global brain asymmetry (particularly for the thickness model and the volume model) are merely contributed by a more general cortical atrophy that occurs with aging. We randomly sampled the brain regions for each “hemisphere” group and conducted the same predictive modeling. That is, a set of regions were randomly selected as features of one “hemisphere” (i.e., simulated “left hemisphere”), and the remaining regions as features of the other “hemisphere” (i.e., simulated “right hemisphere”). This process was repeated for 1000 times, and we found that these random models achieved perfect prediction accuracies (100%) for both thickness and volume. Such high performance was mainly contributed by the increased “hemispheric differences” in the random sampling data (i.e., differences between any two brain regions) which is much larger than the actual differences between the homologous regions. Moreover, when applying these random models to the aging dataset, we obtained various prediction performance for both cortical thickness (from 50% to 100%, mean=78.10%, std=19.14%) and volume (from 50% to 100%, mean=72.59%, std=23.28%). More importantly, the prediction performance showed correlations of different strengths with aging (cortical thickness: from -0.99 to 0.99, mean=-0.12, std=0.78; volume: from -0.99 to 0.99, mean=-0.12, std=0.78). These various results with random sampling data suggest that general cortical atrophy that occurs with aging does not guarantee the specific aging-related decrease observed in the global brain asymmetry. Moreover, these results further supported the specific significance of hemispheric differences in aging.

### Deviations from the typical brain asymmetry in aging-related diseases

The pronounced age-related changes in the global brain asymmetry observed above led us to hypothesize that individuals with aging brain disorders may exhibit more pronounced deviations from the reference asymmetry patterns seen in their healthy young adult counterparts. In other words, when applying the models trained on the young adults to a patient group, the prediction accuracy might be significantly lower than the matched controls.

Therefore, we applied the young-adult models to patient samples focusing on two types of aging brain disorders: AD and PD to test our hypothesis. In light of the results from the functional annotation analyses, this analysis focused on potential changes of the thickness model prediction in AD and the volume model prediction in PD. We identified 34 with AD, and 97 with PD according to their reports in the UK Biobank (see Materials and Methods). The models trained in the healthy young adults (i.e., HCP) were applied to each patient group (Fig. 4). Most of the AD and PD patients were around 70 years old or even older (Fig. S3), and thus, no enough data were available to chart the aging trajectories for patient groups. Here we focused on group-level comparisons between patients and matched control samples. Prediction results showed that in patients with AD, the accuracy for the cortical thickness model was 47.06% – around chance level – and significantly lower than that in the matched control samples (*p* = 0.007, FDR corrected; Fig. 4A). In patients with PD, the prediction accuracy for the volume model was 77.84% which showed a trend of lower performance compared to the matched controls (*p* = 0.069, uncorrected; Fig. 4F). In contrast, the performance for either the volume model in AD (82.35%) or the cortical thickness model in PD (70.10%) was similar to that in the matched controls (*p*s > 0.40; Fig. 4C, 4D), suggesting that aging-related changes in the global brain asymmetry are disease-dependent. Moreover, we also identified 94 patients with all-cause dementia and 105 patients with all cause Parkinsonism (see Materials and Methods). Results in these groups were similar to those for the two groups above (Fig. S4), except for a trend for an additional difference in the prediction performance for the volume model in all-cause dementia (76.60%, *p* = 0.048, uncorrected). No disease-related differences were found for the surface area model (*p*s > 0.40; Fig. 4B, 4E, S4B, S4E).

**Fig 4.**
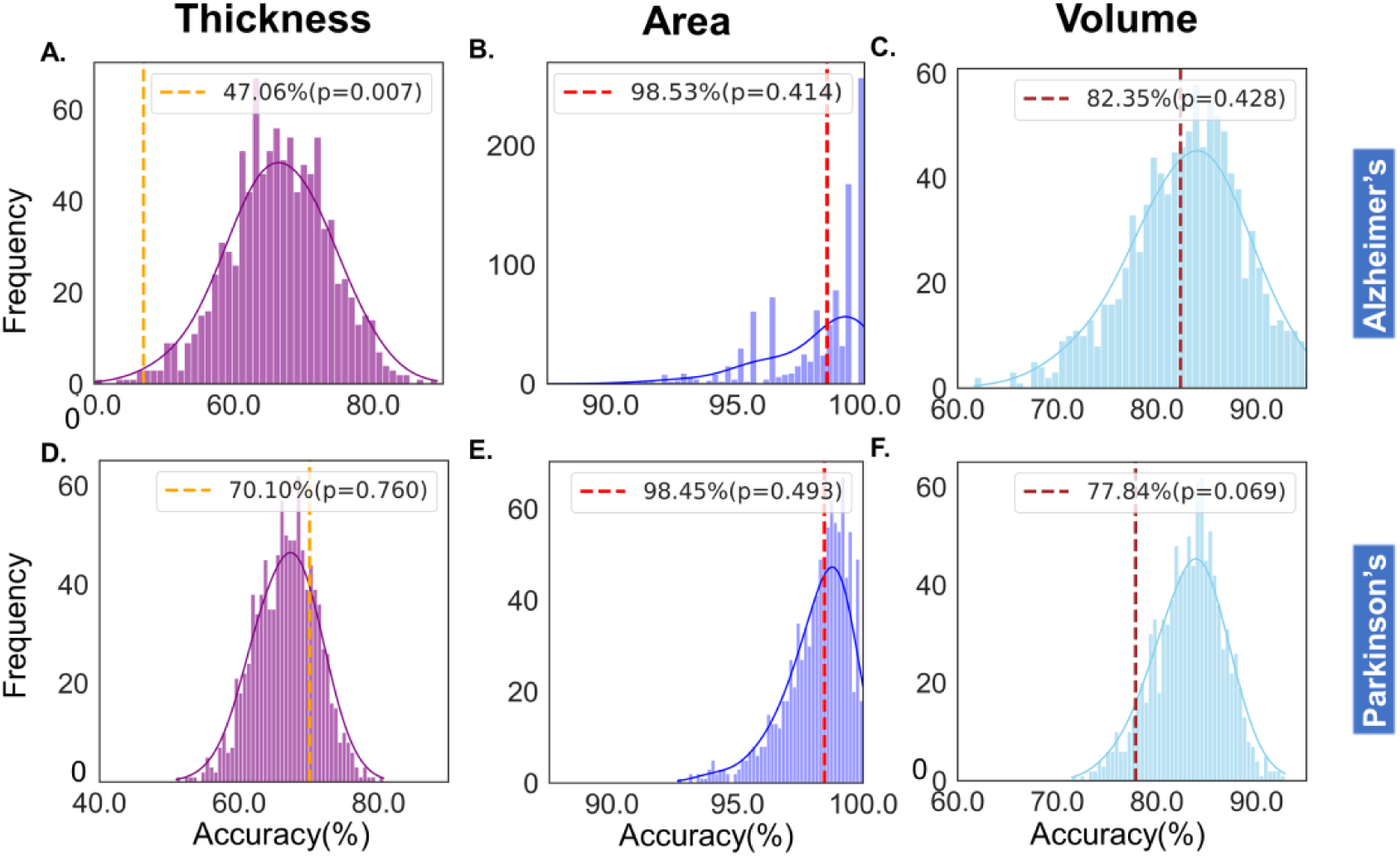
Multivariate brain asymmetry in disorders of brain aging. We mainly focused on the potential changes of the thickness model prediction in AD and the volume model prediction in PD, and also included the others as a comparison. The dashed lines indicate the classification accuracies in the patient groups, while the histograms demonstrate the accuracy distribution in the matched control samples.

Taken together, our results established a clear link between the global brain asymmetry and healthy aging, and showed differential deviations of the brain asymmetry in different aging-related disorders. These results at the population level provide further evidence for the functional relevance of the multivariate brain asymmetry. The proposed models also offer a promising approach for assessing healthy aging, and the phenomenon of “left-hemi aging” provides new insights for a better understanding of aging and related diseases.

### Global brain asymmetry and risk genetic factors

To further study the phenomenon of the deviation in the multivariate brain asymmetry in the aging-related diseases, we explored the effects of relevant genetic risks on the multivariate brain asymmetry in aging. In this analysis, we focused on the APOE gene which have been identified as the most prevalent common genetic risk factor for AD and PD^35,46^. We included individuals who had not been diagnosed with AD or PD but carried various APOE genotypes, focusing on two subsets of samples: the homozygous ε2 carriers (ε2ε2; N = 194) and homozygous ε4 carriers (ε4ε4; N = 789), given that the ε2 and ε4 alleles are the most prominent protective and risk factors, respectively^47,48^.

No differences in age (*t* = 1.30, *p* = 0.19) or sex (*t* = 1.68, *p* = 0.092) were found between the ε2ε2 and ε4ε4 samples. We also did not find any differences in the overall prediction accuracies of either of the three models (thickness: *t* = -1.30, *p* = 0.20; volume: *t* = -0.69, *p* = 0.49). Interestingly, we found that the two genotype groups showed distinct aging patterns in the results for the volume model (Fig. 5C). Specifically, the ε4ε4 samples showed a significant decrease in the classification accuracy of the volume model (*r* = -0.56, *p* < 0.001, corrected), while the ε2ε2 samples showed a significant increase (*r* = 0.61, *p* < 0.001, corrected). In terms of the thickness model’s accuracy, the ε2ε2 carriers showed an increasing trend before the age of 60 (*r* = 0.55, *p* = 0.063), while the ε4ε4 carriers seems to be more stable across aging (*r* = 0.17, *p* = 0.60) (Fig. 5A). The correlation in the ε2ε2 carriers was significantly higher than in random samples with other genotypes, as revealed in the random resampling analysis (*p* = 0.0001) (Fig. S6). This suggests a potential protective effect of ε2ε2 genotype regarding the global pattern of brain asymmetry in aging. No clear differences were found in the surface area model results (Fig. 5B). Please refer to Fig. S5 for results of other AOPE genotype samples (i.e., ε2ε3 ε3ε3 ε3ε4), which mostly showed intermediate effects between the two extreme groups.

**Fig 5.**
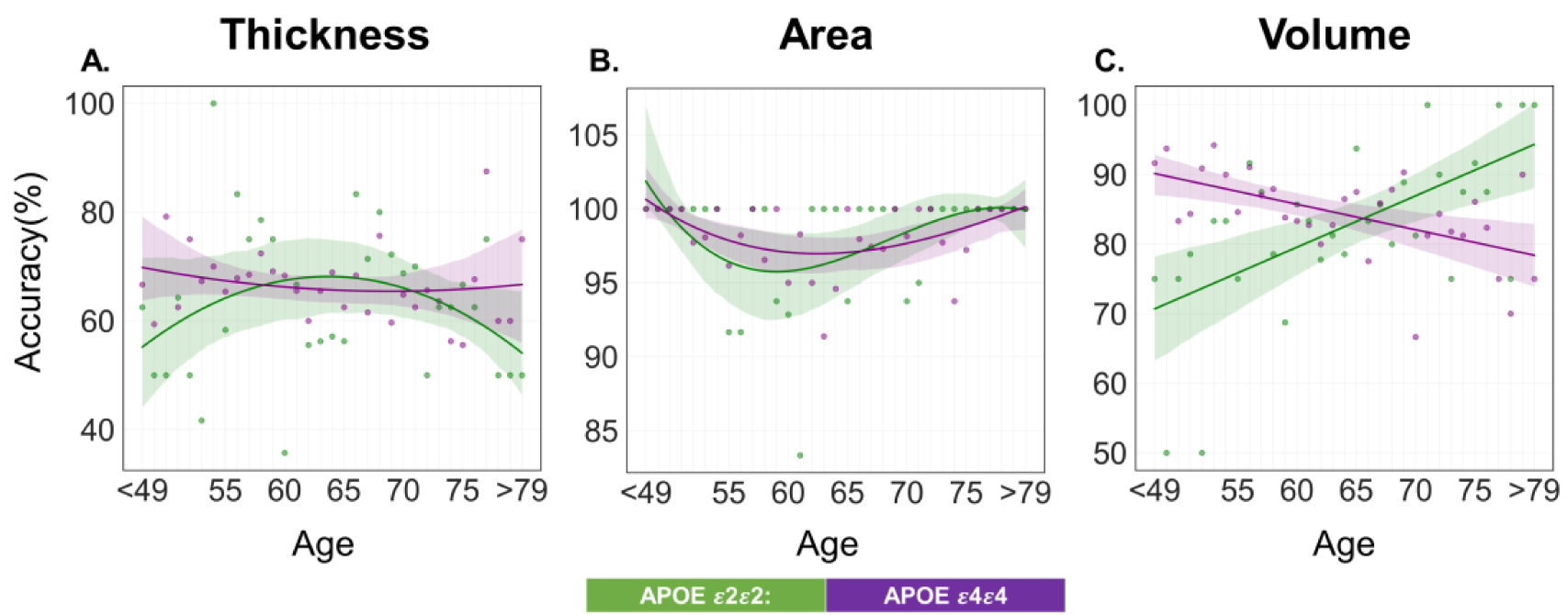
Multivariate brain asymmetry and AD-risk genetic factors. APOE ε2 is typically considered to be a protective factor for late-onset AD and APOE ε4 has been considered as a risk factor. Note that, considering the limited sample size of each age year of below 49 years old in the dataset, samples with an age of younger than 49 years old were grouped together when calculating the statistics (e.g., prediction accuracy). Similarly, samples of above 79 years old were also grouped.

In summary, the above results shed light on a connection between the multivariate brain asymmetry and the genetic risk of aging-related diseases.

### The global asymmetry index and its phenotypic associations

To establish a connection between global brain asymmetry and individual differences in other phenotypes, we introduced an individual-level index termed “global asymmetry phenotype (GAP)” (see Methods). Briefly, the GAP index was defined as the number of correct predictions of the two hemispheres for one individual.

For the thickness and volume model, the association analyses of the GAP index showed significant associations with age^2^ (*p* = 6.77e-4) and age (*p* = 7.02e-8), respectively, aligning with curve fitting results depicting group-level aging trajectories of the prediction accuracy. Additionally, the thickness and volume GAP index demonstrated significant associations with the general cognitive capacity (thickness: *p* = 4.34e-4; volume: *p* = 4.79e-7) and handedness (right-handers > left-handers: *ps* < 1e-100). Intriguingly, we found no significant associations for the surface area model between the GAP index and either variables related to age or cognitive functions, indicating specificity in the observed associations for the thickness and volume model. In addition, we found that in-scanner head motion as one of the major sources of artifact in MRI showed inconsistent correlations with the GAP index of the three models: while the correlations for the thickness model (*t* = -4.71, *p* = 2.54e-6) and the volume model (*t* = 4.29, *p* = 1.82e-5) were in opposite directions, the correlation for the surface area was not significant (*p* = 0.32). The mixed findings indicate that although head motion is somewhat correlated with both aging and the global brain asymmetry, it is unlikely to be a dominant factor in accounting for the observed changes in the global brain asymmetry with aging. The observed associations with age and cognitive functions remained significant when including additional scanning-related variables including head motion as confounding variables in the regression models (see Materials and Methods).

Finally, the phenome-wide association analysis revealed further significant effects, encompassing various brain imaging variables and other factors linked to health (e.g., cardio-respiratory fitness and number of self-reported non-cancer illnesses) and handedness (e.g., birth weight and maternal smoking) (Fig. S7; *p* < 4.64e-6, Bonferroni corrected). Please refer to Supplemental Dataset S4-S6 for additional details. Together, these results collectively underscore the functional relevance of the global brain asymmetry at individual level.

## Discussion

Understanding the structural basis of the cardinal lateralization in the human brain is a fundamental but unresolved question in human neuroscience. To fill in this gap, in the present study, we leveraged cutting-edge machine learning techniques to model the hemispheric differences in brain strucutre, and investigated their functional relevance through a series of large-scale analyses. Our study, from a multivariate perspective, unveiled previously unidentified hemispheric differences in the brain, and demonstrated that the global brain asymmetry was associated with various functions including hand motor and emotion processing. We also showed divergent aging trajectories of the global asymmetry in different morphology which constitutes an important characteristic feature of brain aging (we termed the “left-hemi aging” model). We further revealed differential deviations of the brain asymmetry in different aging-related diseases including AD and PD, and demonstrated distinct aging patterns in the brain asymmetry in individuals with different APOE genetypes. Finally, we introduced the individual-level GAP index and revealed significant associations between GAP and age, as well as individual differences in various phenotypes including cognitive functions, handedness, regional brain measures, and health-related variables. Overall, these results established a link between structural asymmetry, functional lateralization, and brain aging in health and disease. The present study proposed a novel multivariate approach for modelling the inter-hemispheric differences in the human brain. This approach stands apart from that used in existing studies which typically focused on examining differences of homologous regions and reported inconsistent results concerning the associations between brain structural asymmetry and functional lateralization^1,49^. These inconsistent results might stem from the methodological limitation that overlooked complex patterns of the hemispheric differences. We reasoned that the functional hemispheric differences should manifest in global patterns rather than merely the constituent local differences in structure^3,16,22^, and that the global differences should better interpret the functional lateralization in the human brain. As expected, our multivariate modeling results of all the three morphological features showed high classification performance (cortical thickness: accuracy = 87.75%; surface area: 100%; and volume: 99.02%) as well as good reproducibility in independent samples. These results are in line with a recent study which applied a lasso-based linear regression approach but only focused volumetry differences in a much smaller sample size (*N* = ∼200)^50^. Moreover, the functional annotation analyses showed that the revealed hemispheric differences are closely associated with lateralized functions in the brain, including hand movement (left dominance^51^), attention (right dominance^52^), and emotion procesising (right dominance^14^). The association analyses of the individual-level GAP index showed supportive data for the links with handedness and cognitive functions. These results provide supporting evidence for the functional relevance of the proposed global brain asymmetry. The modelling results complement prior observations of minimal left-right differences in brain structure^1,49^ and weak correlation between structural and functional asymmetries^16,53^, and offer an important step towards understanding the structural basis of functional lateralization. Future studies are merited to establish the causal relationship of the structure-function links.

The distinct functional correlates of the three morphological metrics (i.e., cortical thickness, surface area, and volume) were in line with previous findings that different morphological measures are evolutionarily, genetically, and developmentally distinct (e.g., ^1,54,55^). In addition, we noted that the surface area model achieved a relatively higher classification performance than the thickness model, which echoes previous findings that surface area usually showed relativley larger hemispheric differences^1^. These differences may, to an extent, be driven by a potential bias in the parcellation scheme, as the surface-based parcellation used in the present study is more likely to introduce such bias in the sizes of parcellated regions (i.e., surface area). Specifically, larger regions within the parcellation atlas may more likely result in larger size in individual data. In contrast, cortical thickness measures regional distance between the pial surface boundary and the gray-white matter boundary (i.e., along an axis perpendicular to the cortical surface), and thus it depends more on the gray-white matter contrast of individual brain images and would be less likely biased by the parcellation scheme. The potential impact of parcellation dependence will be an important consideration for future studies. Nonetheless, this confunding effect does not undermine the core findings of this study, primarily derived from the thickness and volume models.

We further revealed that the global brain asymmetry changes with aging, by showing divergent aging trajectories of the model prediction of different morphology. Specifically, prediction accuracy results for the volume model exhibited a continuous linear decrease from age 50 to 80, indicating apparently on-going deviations of the multivariate hemispheric differences in aging brains compared to young adults. The thickness model showed a decrease after 65 years old, while the surface area model displayed a relatively stable aging trajectory. Interestingly, we found that the turning point of 65 years old in the results of the thickness model aligns with the age after which AD becomes increasingly prevalent. Patients with AD starting before 65 years old (early-onset AD) and after 65 years old (late-onset AD) show different clinical characteristics^56^. The results indicating that the two hemispheres become less distinguishable with aging are in line with a recent study showing that the asymmetries in the hemisphere-specific brain age were smaller at a higher age^57^. These results may reflect changes in a compensatory function or a de-differentiation process of the two hemispheres in brain aging, which may underlie aging effects in various functions. In addition, it is unlikely that the observed aging-related changes in the global asymmetry is simply a result of more general cortical atrophy that occurs with aging. On the one hand, the prediction models for three metrics did not show identical aging trajectories, suggesting metric-specific effects of aging. On the other hand, the predictive modeling of simulated “hemispheric” data showed various aging-related changes in the prediction performance, which largely contrasted with the aging changes in the global brain asymmetry. Thus, we believe that the aging-related changes in the global brain asymmetry is specific to the hemispheric differences in aging.

Moreover, we found that these changes observed in the volume and thickness models were primarily attributed to an increased misclassification of left hemispheres as right hemisphere with aging. This suggest that the left hemispheres become less distinguishable from the right hemisphere as individuals age. These findings challenge the traditional “right hemi-aging” model, which posits greater age-related decline in the right hemisphere than the left^27^. Note that the “right hemi-aging” model was proposed mainly based on behavioural observations. Such an extension from behaviours to brain organization could not be justified given the not yet established link between them. Our findings align more with the HAROLD model which indicates less lateralized functions with aging in the brain^28^. Notably, while the HAROLD model only describes local differences between homologous regions, the left-hemisphere-linked changes observed in our study reflect a global shift across the entire brain. This leads us to term this phenomenon “left-hemi aging”. We additionally speculated that these left-hemisphere-linked changes could underlie age-related changes in left-lateralized functions such as language lateralization ^28,58^. Although this new model provides fresh insights into healthy brain aging and aging-related diseases, the underlying mechanisms remain yet to be fully elucidated. The human brain exhibits left-right asymmetry, a phenomenon evident early in development^59^. Gene expression and epigenetic regulation differ between hemispheres, potentially driving asymmetries in cellular distributions, connectivity, neurotransmitters, and protein expression^26^, which may further contribute to vulnerabilities in one hemisphere. Further research is imperative for a more comprehensive understanding of the underlying mechanisms.

In comparison with related studies, two recent investigations explored lifespan brain asymmetry using a multivariate framework. Saltoun and colleagues employed singular value decomposition (SVD) on the 85 brain regional measures of the volume asymmetry index, revealing dissociable asymmetry patterns with distinct phenotypic associations^60^. Similarly, Roe and colleagues utilized a clustering algorithm on asymmetry age trajectories, identifying three clusters with unique asymmetric thinning properties across the adult lifespan^31^. Both studies, consistent with our findings, presented supportive data for brain asymmetry changes in aging. For instance, Saltoun et al. (2023) revealed notable age-related changes in the expression of all asymmetry patterns except the first one. Roe et al. (2021) reported accelerated thinning in the previously thicker homotopic hemisphere during aging and in patients with AD. However, their primary focus was on decomposing brain asymmetry into relatively independent spatial patterns. This was based on the inter-regional similarity in either the individual differences of asymmetry measures or the asymmetry age trajectories. In essence, an unsupervised approach guided both studies. In contrast, our work adopted a supervised approach for investigating inter-hemispheric differences. Specifically, we trained multivariate prediction models to explicitly classify the two hemispheres, enabling the identification of the left-hemi aging phenomenon. Moreover, these trained models can be easily applied to new datasets, facilitating the study of global brain asymmetry in brain disorders and other related individual variations.

Besides the aging-reletad changes, we further revealed diagnosis-specific deviations of the global brain asymmetry in two aging-related diseases (AD and PD). The most prominent association was a link between the thickness model and AD. Specifically, we found that patients with AD showed a chance-level classification accuracy (i.e., 47%) in the cortical thickness modeling results, which was significantly lower than that in matched controls. Similar effects were found when including other forms of dementia (i.e., 56%). These results suggest an accelerated reduction of the multivaraite hemispheric differences in cortical thickness in AD. This pattern of results is in line with recent findings of accelerated asymmetric thinning of homologous regions in AD with longitudinal cohorts^31^. Moreover, our analyses with PD patients showed a similar effect as AD but in the volume model (i.e., 77.62%). It is important to note that the PD-related effects did not survive multiple testing correction, and independent replication is needed. Nonetheless, these differential effects in AD and PD echo the observations in the functional annotation analyses: AD showed the strongest correlation with the thickness model while PD showed the strongest correlation with the volume model. In addition, we noted that the functional annotation analyses also revealed strong correlations with anxiety disorders and psychiatric disorders. Thus, we speculate that patients with anxiety or depressive disorders may show alterations in the global brain asymmetry. While little differences were found in regional asymmetry of brain morphology^12^, such a new hypothesis from a multivariate perspective merits further investigation, e.g., via the ENIGMA consortium^11,61^. Taken together, the links between the multivariate brain asymmetry and aging and aging-related diseases have reinforced the functional relevance of the proposed multivariate hemispheric differences. Furthermore, the proposed multivariate approach stands as a promising tool for the assessment of healthy brain aging, and presents significant opportunities to deepen our understanding of the aging process and its associations with related diseases.

Lastly, we found that individuals who were not diagnosed as either AD or PD but had a higher genetic risk for such neurodegenerative disorders (i.e., APOE ε4) showed differential aging trajectories in the model performance, compared to those with a protective variant (i.e., APOE ε2). These differences were particularly apparent in the volume modeling results. The observed effects of APOE genotypes should not be attributed to other confunding factors such as different scanners and population, as samples of both the two groups (i.e., APOE ε2ε2 and ε4ε4) were from the the same dataset, and no differences in age or sex were found. These results provide valuable insights into the connections between the global brain asymmetry and the genetic risk of aging-related diseases. This discovery introduces a new candidate for understanding the changes in brain asymmetry during the aging process.

There are limitations to this study. First, while substantial evidence from different perspectives in the present study supports the connections between global brain asymmetry, healthy aging, and cognitive functions, it remains unclear whether these findings can be replicated in independent datasets or if the prediction models for the hemispheric differences are applicable to other disorders, such as autism spectrum disorder^62^, obsessive-compulsive disorder^9^ and major depression^11,12^. Further studies are necessary to demonstrate the effectiveness of the proposed approach in other cohorts. Secondly, this study primarily addressed the cortical measures of the brain, including cortical thickness, surface area and volume. Existing research has highlighted subcortical asymmetry and associated alterations observed in various brain disorders. Future investigations should explore aging trajectories of subcortical asymmetry, encompassing the examination of subcortical nuclei and ventricles^11,63,64^. Furthermore, while the permutation-based approach employed for estimating feature importance offered valuable insights into the trained models at group level, newer explanation techniques such as SHapley Additive exPlanations (SHAP) and Local Interpretable Model-agnostic Explanations (LIME) could more precisely indicate which features influenced individual predictions^65^.

In sum, we introduced a multivariate approach for modeling and understanding the inter-hemispheric differences by integrating advanced machine learning techniques, and presented the largest-ever analysis of brain structural asymmetry in relation to cognition, aging and aging-related diseases. Our proposed multivariate approach opens a new avenue for understanding the hemispheric specialization in the human brain and bridges the long-existing gap between the subtle hemispheric differences in brain structure and the obvious functional lateralization. Furthermore, our study introduces a novel method for assessing both healthy and diseased brain aging, making a significant contribution to our understanding of aging-related diseases.

## Materials and Methods

### Datasets

Brain imaging data from three large-scale datasets, i.e., the HCP, UK Biobank, and a large-scale neuroimaging meta-analytic database were used in the present studies.

#### HCP

The HCP provides high-quality brain MRI data from 1,113 young adults (606 females, age range: 22–37 years, at the time of scanning) (https://humanconnectome.org/). The dataset is publicly available and was approved by the Institutional Review Boards associated with HCP. FreeSurfer-derived morphometric measures^66^ including regional cortical thickness, surface area, and volume of the 68 cortical regions (34 per hemisphere) in the Desikan-Killiany atlas^67^ were extracted for each individual. HCP has established well-designed quality control protocols for MRI data collection and image processing^68^, and MRI data labeled as “unusable” were not used in the present work. This high-quality dataset was used to demonstrate the multivariate brain asymmetry approach and establish reference models for making predictions in older cohorts. Only samples with data of all the 34 regions were included for model training and validation. Furthermore, Cohen’s d effect-size contract maps for all HCP fMRI tasks were acquired through ConnectomeDB^69^. We utilized all 86 contract maps (Cohen’s d) related to seven functions including working memory, gambling, motor, language, social cognition, relational processing, and emotion processing. For each contract, brain maps of two smoothing levels (2mm and 4mm) were available in the HCP. These maps form an integral part of the brain functional data used for functional annotation of feature importance, as described below.

#### UK Biobank

UK Biobank is a general aging population cohort (N = 502,411; median age = 64 years; range 44-83 years). The data collection has been described elsewhere^70–72^. Informed consent was obtained for all participants, and the UK Biobank received ethical approval to disseminate the data publicly. For this study, we used data obtained as part of research application 75807, with Dr. Xiang-Zhen Kong as the principal applicant. A subset of participants was included in this study for whom brain imaging data were available – in total, this subset had 43,102 participants. The median age of these participants was 64 years (range: 44-82 years), and 22,708 subjects were female (52.68%). Brain imaging features were derived from the cortical parcellation using the Desikan-Killiany atlas from FreeSurfer. These features were cortical thickness, surface area (of white and pial surface), and volume of each of the parcellations in the atlas. We calculated surface area by averaging the measures of the white and pial surface. All of the brain imaging features used in the analyses were derived, validated and made available from the neuroimage processing pipeline of the UK Biobank^72,73^. Measures of all three metrics of the temporal pole and the surface area of the insular were excluded due to limited data quality. Visual inspection of each of the remaining measures showed a nearly perfect normal distribution. Only samples with complete data for all remaining parcellations were included for the prediction analyses. This resulted 42913 participants with no history of age-related disorders (see below), enabling the analysis of typical trajectories of the global asymmetry in healthy aging. It’s worth noting that the data available for this study had already undergone extensive quality control (QC), which included automated classification (based on a large-number of QC-specific merics) and manual reviews^72,75^. Thus, we did not exclude additional participants from our current analyses based on these QC metrics. Instead, when applicable, we included the QC metrics as potential confounding variables in our regression models (see *Phenotopic associations of multivariate brain asymmetry at individual level*). For detailed information on image acquisition protocols, quality control, analysis pipelines, and derived measures, please refer to Resource 1977 of the UK Biobank (biobank.ndph.ox.ac.uk/ukb/ukb/docs/brain_mri.pdf). Individuals with various age-related brain disorders were identified using variables with these Field IDs: 42018 for “all cause dementia report”, 42020 for “Alzheimer’s disease report”, 42030 for “all cause parkinsonism report”, and 42032 for “Parkinson’s disease report”. We also made use of genotype data for two single-nucleotide polymorphisms (SNPs) of the APOE gene (i.e., rs7412, and rs429358) which are well-known as the most prevalent genetic risk factor for late-onset AD. These two SNPs determine the genotype of APOE. The sample sizes for each genotype subgroup were as follows: N = 194 for homozygous ε2 carriers; N = 4,433 for ε2ε3 carriers; N = 21,273 for homozygous ε3 carriers; N = 8,294 for ε3ε4 carriers; and N = 789 for homozygous ε4 carriers. Detailed sample sizes for each group are shown in Fig. S3.

#### NeuroSynth

Neurosynth is a large-scale brain activation coordinate database of published neuroimaging studies^76^. It uses text-mining techniques to detect frequently used terms as proxies for functional concepts of interest in thousands of papers from the neuroimaging literature. Terms that occur at a high frequency in a given study are associated with all activation coordinates reported in that publication, allowing for an automated term-based meta-analysis.

The latest version of this dataset was used (version 0.70), which includes over 500,000 activations from 14,371 studies (https://neurosynth.org/). This database was used for functional annotation of the reference prediction models.

### Multivariate brain asymmetry analyses in young adults

#### Model training

Unlike prior brain asymmetry studies that directly compared features of homologous regions in the two hemispheres^1,11^, the present study introduced a novel multivariate framework. In the framework, we treat all regional measures within one hemisphere as a collective entity, aiming to test whether and to what extent, the morphometric features of these regions can distinguish the left and right hemispheres. We demonstrated the framework using the FreeSurfer-derived morphological data from a set of young adults in the HCP (see *Datasets*). Specifically, we used morphological data (i.e., regional cortical thickness, surface area, and volume) of the 34 regions in each hemisphere as features, and trained an ensamble machine learning model for classifying the left and right hemispheres. The pairwise correlations between the 34 regions remained predominantly below 0.70 for all three morphological metrics, suggesting absence of collinearity between these features (Fig. S8). An ensemble model can consist of various branches/submodels, employing diverse machine learning algorithms such as LDA (Linear Discriminant Analysis), MLP (Multilayer Perceptron), SVM (Support Vector Machine), decision tree, and random forest. Each of these contributes to the overall ensemble model, enabling it to extract more detailed information from the original data (Fig. S1). Normalization and feature decomposition were also integral components of the ensemble model. We employed an automated machine learning toolkit *auto-sklearn*^34^, available at https://automl.github.io/auto-sklearn/. This toolkit harnesses recent advancements in Bayesian optimization, meta-learning, and ensemble construction, liberating machine learning applications from the challenges of algorithm selection and hyperparameter tuning ^34^. As the HCP dataset has more females (*N* = 606) than males (*N* = 507), in order to ensure the balance between the sex groups and avoid potential bias in the trained models, we first randomly removed 99 female participants. The final dataset consisted of 1014 participants, half male and half female. When training the multivariate models, the train_test_split function from *sklearn* was used for splitting samples into random train and test subsets. In the present work, we sampled a training set of 70% of participants (male-female balanced), and held out the remaining 30% for testing our classifiers. The classification accuracy was computed within the testing set to evaluate the overall performance of the trained model. We conducted repeated the train-test sampling procedure, consistently yielding similar model performance. Subsequent analyses were conducted using initial set of models that we trained, separately for cortical thickness, surface area, and volume. We also checked the single branch with the highest ensemble weight within each of the models.

#### Feature importance estimation

To rank the features in terms of their contribution to the prediction models (i.e., feature importance), we used a permutation-based approach. The fundamental concept of this approach is that a variable is deemed important if its exclusion adversely affects overall prediction accuracy. This empirical approach has been well established and is described below:

*Inputs: fitted predictive model **m**, and tabular dataset **D***

*First, compute the reference performance score **s** of the model m on data **D** (in the present study, the classification accuracy)*.

*For each feature **j** (i.e., column of **D**): For each repetition **k** in 1,…,**K**:*

*Randomly shuffle column **j** of dataset **D** to generate a corrupted dataset, **D̃_k,j_***

*Compute the score **s_k,j_** of model **m** on the corrupted data, **D̃_k,j_***

*Compute importance **i_j_** for feature **j** defined as:*

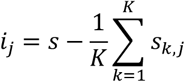

Thus, we obtained an importance score for each of the cortical regions in each prediction model. A higher importance score indicates a greater contribution to the hemispheric classification. Please note that this procedure was primarily for ranking the features for the subsequent functional annotation, rather than making inferences about specific individual features. The approach used in this study differs from the potentially problematic practice discussed by Bzdok et al. (2020), where Lasso (least absolute shrinkage and selection operator) model coefficients or the coefficient *p* values of simple linear models were used for statistical inference^40^.

#### Functional annotation of feature importance

We used the large-scale neuroimaging database, NeuroSynth, to provide an exploratory functional interpretation of the most important features in the predictive models. Specifically, first we ran a coordinate-based meta-analysis for each term (e.g., “language” and “Alzheimer”) using the *MetaAnalysis* function implemented in NeuroSynth, and obtained a synthetic brain map that suggests an association with that functional concept of interest. The generated maps were in the standard MNI space. To further make the meta-analytic data comparable with the importance scores, we further projected the maps into FreeSurfer’s fsaverage space. The well-validated script *CBIG_RF_projectMNI2fsaverage.sh* from CBIG was used (https://github.com/ThomasYeoLab/CBIG). Next, we used the *mri_segstats* application from FreeSurfer to calculate the average values for each of the 68 cortical regions (34 per hemisphere). Finally, the absolute differences between homologous regions in the two brain hemispheres were calculated, and further tested for correlation with the importance scores estimated for each prediction model. Spearman’s rank correlation was used, along with a spatial spin permutation test (n=10,000) for assessing the nonparametric statistical significance^77^. We repeated these steps above for each of the terms of interest. Two sets of terms were particularly of interest in the present study: one with 590 cognitive terms curated in a previous study ^14^, and one with 40 terms related to various brain disorders, including “alzherimer”, “parkinson”, and “depression”. Note that these terms present all available terms in the database after filtering out non-specific terms which is not specific to brain functions (e.g., “ability”, “tasks”, “usually” and “accuracy”) (https://neurosynth.org/analyses/terms/). In other words, this filtering is not because they are irrelevant to brain asymmetry or lateralization, but rather due to these terms being functionally meaningless in relation to brain functions. The false-discovery-rate (FDR) correction procedure (*q*=0.05) was used to correct multiple testing. The Neurosynth-based functional annotation relies on large-scale neuroimaging data, is well-established, and has been widely used for the interpretation of brain maps (e.g., ^42–45^). Moreover, with group average functional brain maps from the HCP dataset, we ran an additional analysis for exploring other potential functional correlates. We utilized all 86 effect-size contract maps (Cohen’s d) derived from real fMRI data of a battery of 7 tasks, including working memory, gambling, motor, language, social cognition, relational processing, and emotion processing. A detailed description of these tasks can be found in Barch et al. (2013). The results remained the same for both smoothing levels of the brain maps (i.e., 2mm and 4mm), and results with 2mm were reported in the main text.

### Multivariate asymmetry in the aging brain

Next, we applied the models – already trained on the young adults – to a large cohort of aging adults in the UK Biobank. The aim was to chart the aging trajectories of the global brain asymmetry. Cortical disorganization is known to occur with aging (e.g., ^78^). We hypothesized that this disorganization during aging could be asymmetrical between brain hemispheres, leading us to anticipate varying accuracy of the hemisphere classification models across different age groups. Specifically, for each morphological metric, we trained a reference predictive model as above for classifying the two hemispheres in the HCP cohort. Given the lack of morphological measures for two regions, i.e., the insular (for surface area) and temporal pole (for all three metrics), in the UK Biobank, we retrained the reference models with data from the remaining 32 or 33 regions in the HCP, which resulted in a similar performance in the young cohort. Then, we applied these models to classify the hemispheres for each individual in the UK Biobank. The prediction accuracy of different age bins may reflect aging-related changes in asymmetry patterns. That is, we are interested in aging-related changes in the prediction accuracy within the UK Biobank, rather than decreased prediction accuracy per se compared with that in the HCP, which could be due to many other factors including scanner, imaging protocol, and sample population. A curve fitting analysis of the prediction accuracy and ages was conducted to test whether the aging-related changes were linear, quadratic, or cubic. The best fit was determined according to the Akaike information criterion (AIC). AIC is a likelihood-based technique that is based on a trade-off between parsimony and fit. A smaller AIC score indicates a better fit.

To further understand the observed changes, we also calculated the asymmetry index (AI, AI=(L-R)/(L+R)) for each region of each morphological metric in both HCP and UK Biobank, and tested the similarity of the asymmetry index between the older and the younger cohorts. This is an alternative for studying the age-related changes in brain-wide asymmetry patterns.

In addition, we found that, when we applied the trained models from the HCP young adults to the UK Biobank aging samples, significant aging-related changes were observed in the prediction performance (see Results). We further asked whether the observed aging-related changes in the global asymmetry (particularly for the thickness model and the volume model) is simply part of a general cortical disorganization that occurs with aging. To answer this question, we conducted the same predictive modeling but with random sampling of brain regions from both hemispheres. Specifically, we iteratively select random sets of 32 or 33 brain regions (from both the left and right) and train a model to recognize those sets (i.e., a simulated “left hemisphere”) vs. the remaining 32 or 33 regions (i.e., a simulated “right hemisphere”). Next, we applied these random models to the aging dataset and calculated the prediction accuracy for each age group as in the original analysis. We repeated this process for 1000 times. If these random models also exhibited an age-related decrease in performance, our observed changes might simply reflect a broader, more generalized phenomenon of cortical disorganization. Conversely, if the random models did not show this pattern, it would suggest that the aging-related changes we observed were specific to brain asymmetry, which would further support the significance of brain asymmetry in aging.

### Multivariate brain asymmetry in aging brain disorders

We hypothesized that, if the degree of global brain asymmetry correlates with older age, patients with age-related brain disorders may also present greater deviation from the reference asymmetry patterns, that is, lower predictive performance compared to matched controls. We calculated the predictive performance in the patient groups including AD and PD (see *Datasets*). As a comparison, we randomly sampled the same number of age- and sex-matched controls as the patient group from the remaining data, and calculated a prediction accuracy. The sampling was repeated 1,000 times to obtain a distribution of the prediction accuracy in the matched controls for each brain disorder. By comparing the predictive accuracy of a patient group with the control distribution, a significance value can be derived for the effect of each brain disorder. We applied a significance threshold of a corrected false discovery rate (FDR) of 0.05 to account for multiple comparisons (in total 6 tests: 3 models [cortical thickness, surface area, and volume] × 2 diseases [AD and PD]). Considering the results from the functional annotation analyses, in this analysis our focus was on the prediction performance of the thickness model in AD and the prediction performance of the volume model in PD. Additionally, for comparison, we identified all-cause dementia and all-cause parkinsonism in the UK Biobank (see *Datasets*) and examined all the three models (thickness, volume, and surface area) within these different patient groups.

### Multivariate brain asymmetry and AD-risk genetic factors

We further explored the differential effects of APOE genotypes that are the most prevalent common genetic risk factor for AD^48^. The APOE genotypes of APOE2, E3, and E4 were determined by the two SNPs rs7412, and rs429358. As the APOE2 and APOE4 genotypes have widely proven to be protective and risk factors for AD, respectively, in the present study, we mainly focused on the comparisons between these two extreme groups (i.e., ε2ε2 versus ε4ε4) to maximize the effects. Specifically, we tested the thickness model and the volume model in both ε2ε2 and ε4ε4 samples, and a significance threshold of 0.0125 (i.e., 0.05/4) was considered significant. The results of other intermediate genotype groups (i.e., ε2ε3, ε3ε3, and ε3ε4) were also reported. A significance threshold of a corrected false discovery rate (FDR) of 0.05 to account for multiple comparisons.

### The global asymmetry index and its phenotypic associations

To further establish a connection between the global brain asymmetry and individual differences in various phenotypes, we introduced an individual-level index for the global brain asymmetry based on each individual’s prediction results. Specifically, the index, termed “global asymmetry phenotype (GAP)”, was defined as follows: a score of 2 when both hemispheres were correctly labeled in the classification, a score of 1 when only hemisphere was correctly labeled, and a score of 0 when both hemispheres were incorrectly labeled.

Using the GAP index, we explored associations between the global brain asymmetry and age and individual differences in other phenotypes, including cognitive functions, handedness, regional brain measures, and health-related variables. Regarding the age effects, we conducted regression analyses with the GAP index as the dependent variable, and age, age^2^, and age^3^ as the independent variables. For investigating the associations with cognitive funciotns, ten available variables related to cognitive functions were extracted (Field IDs: 399 for “Pairs matching”, 4282 for “Numeric memory”, 6348 and 6350 for “Trial making”, 6373 for “Matrix pattern completion”, 20016 for “Fluid intelligence/reasoning”, 20023 for “Reaction time”, 21004 for “Tower rearranging”, and 23324 for “Symbol digit substitution”). A sample size of 35785 with all the 10 cognitive variables was obtained. Principal component analysis (PCA) was performed on these variables to derive a single variable capturing general cognitive capacity. Data of handedness was extract with the Field ID 1707. Linear regression was employed to examine associations with the GAP index. Scanning-related variables were controlled for in the analyses. These variables included those with field IDs 54 for “UK Biobank assessment centre”, 25734 for “Inverted signal-to-noise ratio in T1”, 24419 for “Measure of head motion in T1 structural image”, and 25756, 25757, 25758, and 25759 for scanner table position variables. Please note that the head motion metric (field ID 24419) was not a direct measure of motion during scanning but was an estimate based on a cross-validated linear regression model. In the regression model, the dependent variable was a manually assessed QC measure of motion in a separate set of T1 images, and the independent variables were a collection of features associated with structural motion and QC. For detailed information on quality control, analysis pipelines, and derived measures, please refer to Resource 1977 of the UK Biobank (biobank.ndph.ox.ac.uk/ukb/ukb/docs/brain_mri.pdf).

Additionally, we utilized extensive phenotypic data in the UK Biobank and the PHEnome Scan Analysis Tool (PHESANT)^79^ to further expore other potential correlates of the GAP index. Age, sex, and scanning-related variables (as mentioned above) were included as confounding factors. Multiple testing in the PHESANT analysis was performed using Bonferroni correction, with a significance threshold determined by dividing 0.05 by the number of tests performed (0.05/10779 = 4.64e-6). It is important to note that while the massive phenotype data included handedness, early life factors, and physical and mental health records, variables related to language or auditory lateralization were unavailable in the UK Biobank.

## Acknowledgments

This research has been conducted using the UK Biobank Resource under application number 75807, with Xiang-Zhen Kong as the principal applicant. Our study made use of imaging-derived phenotypes generated by an image-processing pipeline developed and run on behalf of UK Biobank. We thank the UK Biobank and the HCP for data sharing. Xiang-Zhen Kong was supported by the National Natural Science Foundation of China (32171031), the China Brain Project (2021ZD0200409), the Fundamental Research Funds for the Central Universities (2021XZZX006), and the Information Technology Center, Zhejiang University. PMT is supported by the US NIH and the Alzheimer’s Association. YP and CF was supported by the Max Planck Society (Germany).

## Data availability

For use of UK Biobank data, an application must be made via http://www.ukbiobank.ac.uk/register-apply/. The HCP data are available via https://www.humanconnectome.org/.

## Code availability

All analyses were carried out as described in the methods. Software versions and relevant parameters are included in the corresponding methods sections. Scripts are available from open access repository (e.g., OSF) upon publication.

## Author contributions

HH: data preparation, data analysis, visualization, and preparing the first draft; YP: study conception and design, data analysis, manuscript editing and critical review; YA: editing the manuscript; CF: manuscript editing and critical review; PMT: manuscript editing and critical review; XK: study conception and design, data preparation, data analysis, results interpretation, visualization, preparing the first draft and manuscript editing.

## Supplemental Datasets and Figures

**Dataset S1-S3. Inter-hemispheric differences in regional thickness (S1), surface area (S2), and volume (S3) of homologous regions within each age group.** Cohen’s d was computed by dividing the mean with the standard deviation. Note that, considering the limited sample size of each age year of below 49 years old in the dataset, samples with an age of younger than 49 years old were grouped together when calculating the statistics (e.g., prediction accuracy). Similarly, samples of above 79 years old were also grouped.

**Dataset S4-S6. PheWAS results for the GAP index with the thickness model (S4), surface area model (S5), and volume model (S6).** Only top associations that survived multiple testing correction were reported. For further details on each phenotype, please refer to https://biobank.ctsu.ox.ac.uk/showcase/search.cgi.

**Fig. S1.**
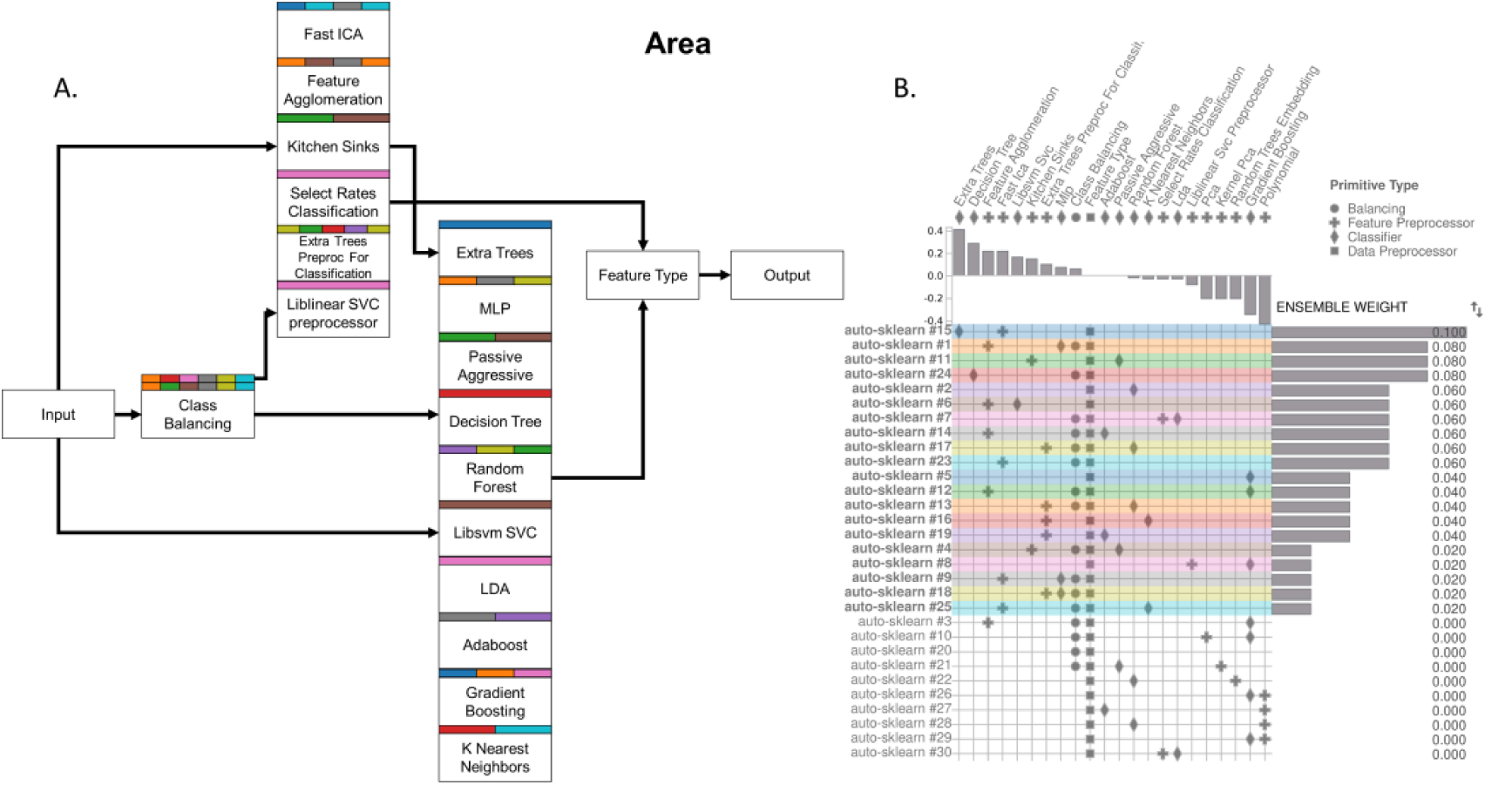
Pipeline matrix for the surface area model. (A) The step-by-step autoML PipelineProfiler flowchart. Header colors code different pipelines. (B) The classifier Extra Trees (i.e., extremely randomized trees) ranked as the top #1. The matrix indicates primitives (e.g., data preprocessing, classifiers, feature preprocessor, and balancing applications; in the *columns*) used by each pipeline (i.e., each branch of the ensemble model; in the *rows*). Bar plots show the primitive contribution (correlation between primitive usage and performance contribution; *top*) and the ensemble weight ranking (*left*).

**Fig. S2.**
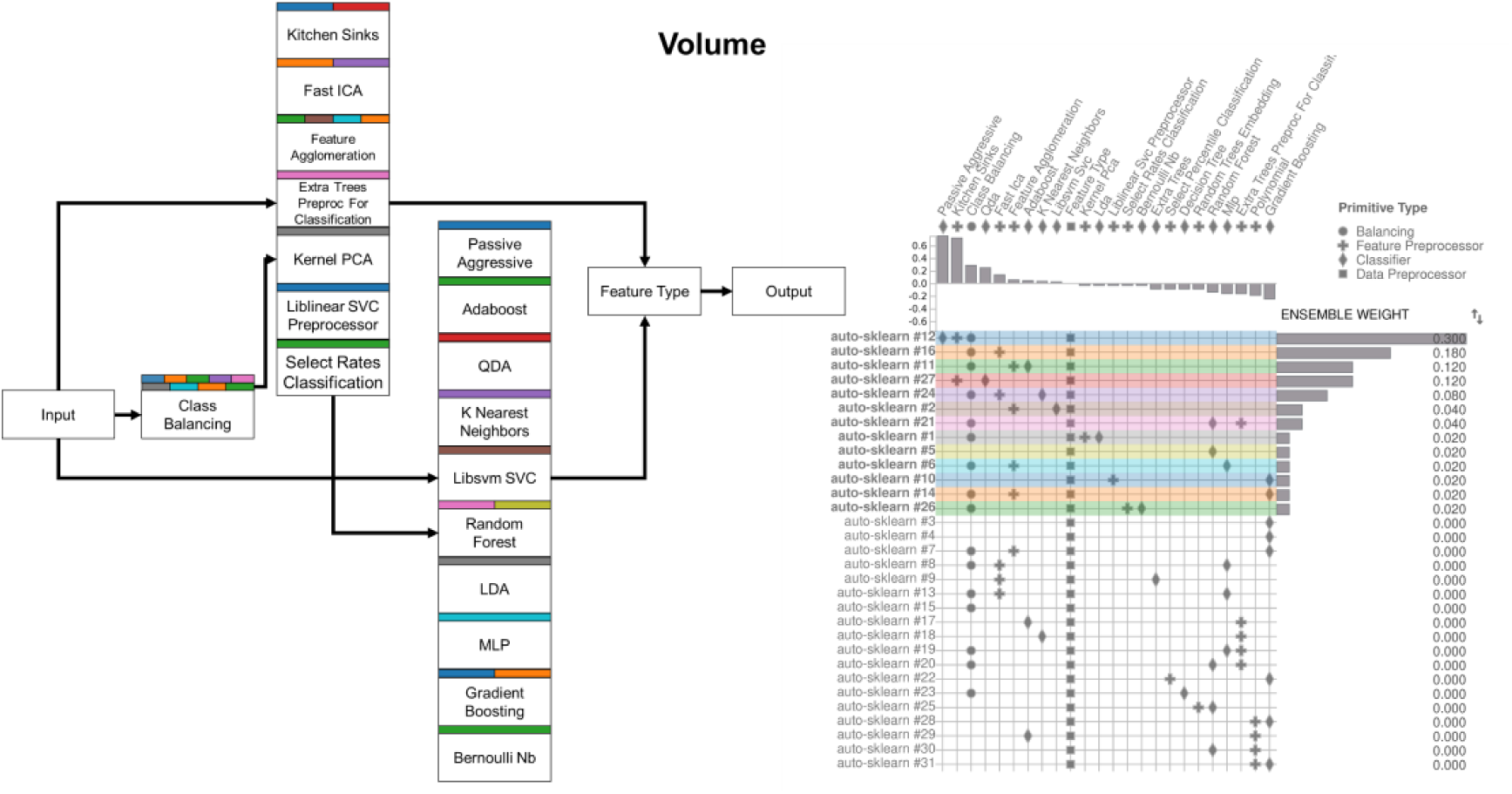
Pipeline matrix for the volume model. (A) The step-by-step autoML PipelineProfiler flowchart. Header colors code different pipelines. (B) The classifier Passive Aggressive ranked as the top #1. The matrix indicates primitives (in the *columns*) used by each pipeline (i.e., each branch of the ensemble model; in the *rows*). Bar plots show the primitive contribution (correlation between primitive usage and performance contribution; *top*) and the ensemble weight ranking (*left*).

**Fig. S3.**
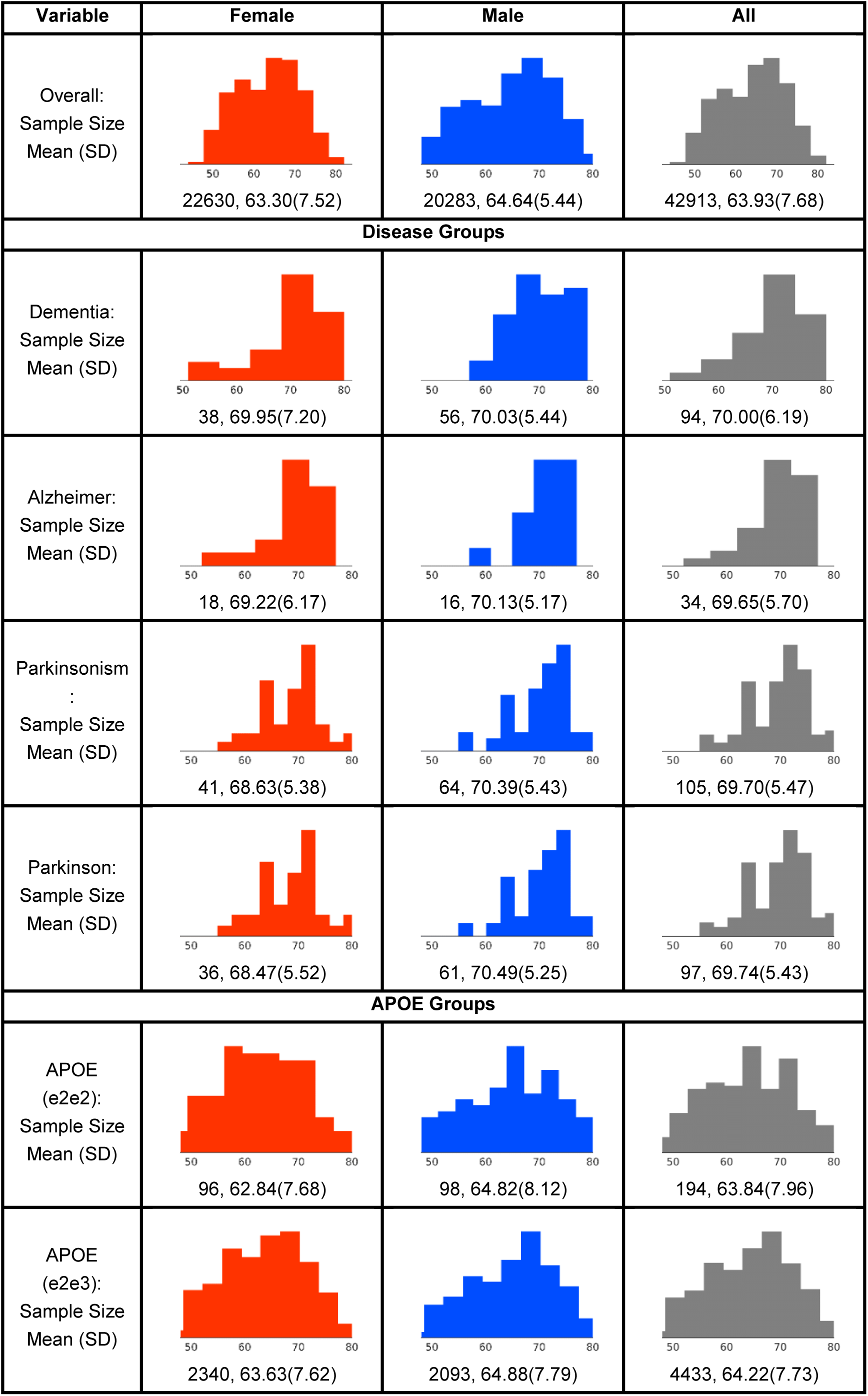
The sizes and age distributions of each group.

**Fig. S4.**
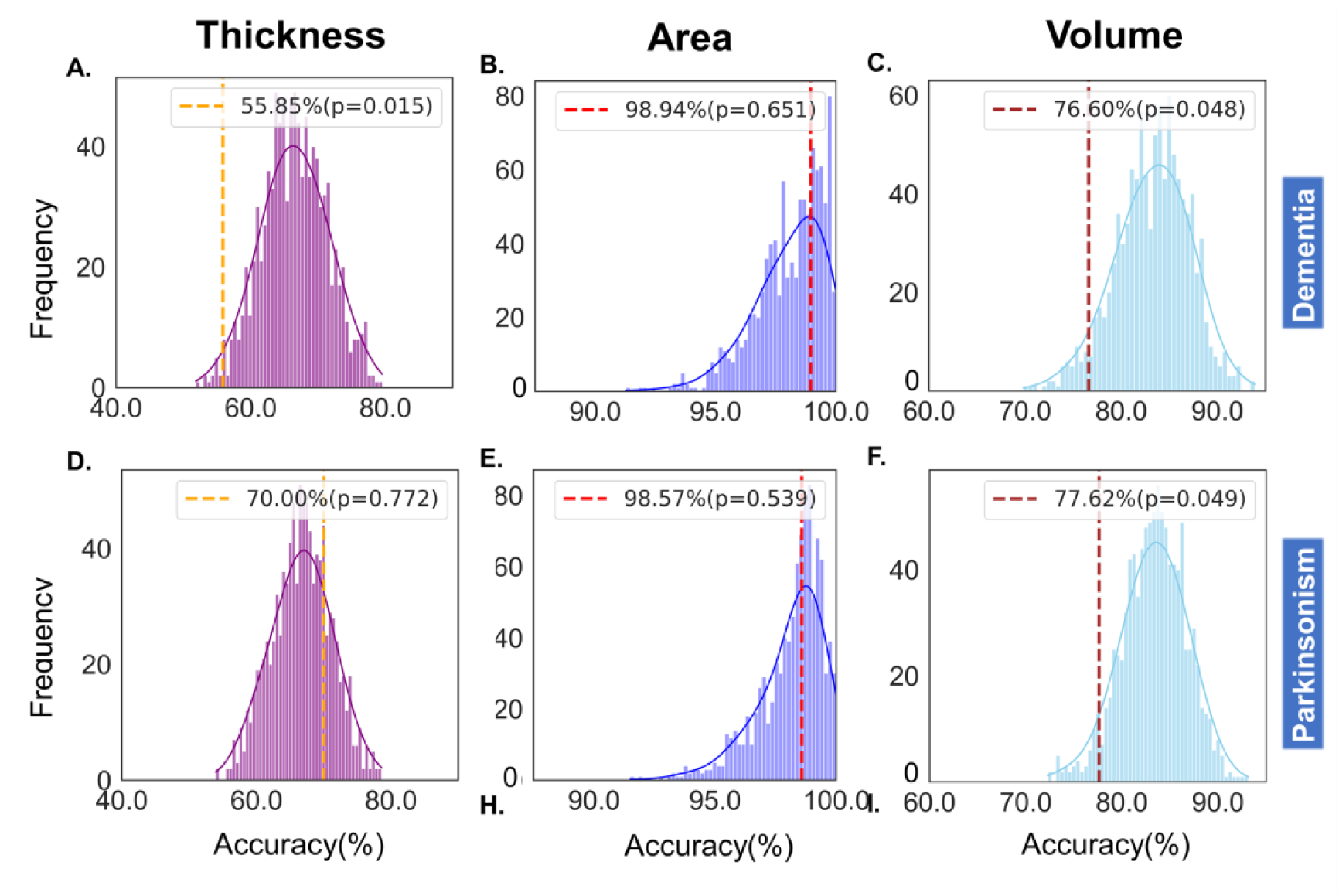
Global brain asymmetry in all-cause dementia and all-cause parkinsonism in the UK Biobank. The dashed lines indicate the classification accuracies in the patient groups, while the histograms demonstrate the accuracy distribution in the matched control samples (repeated for 1000 times).

**Fig. S5.**
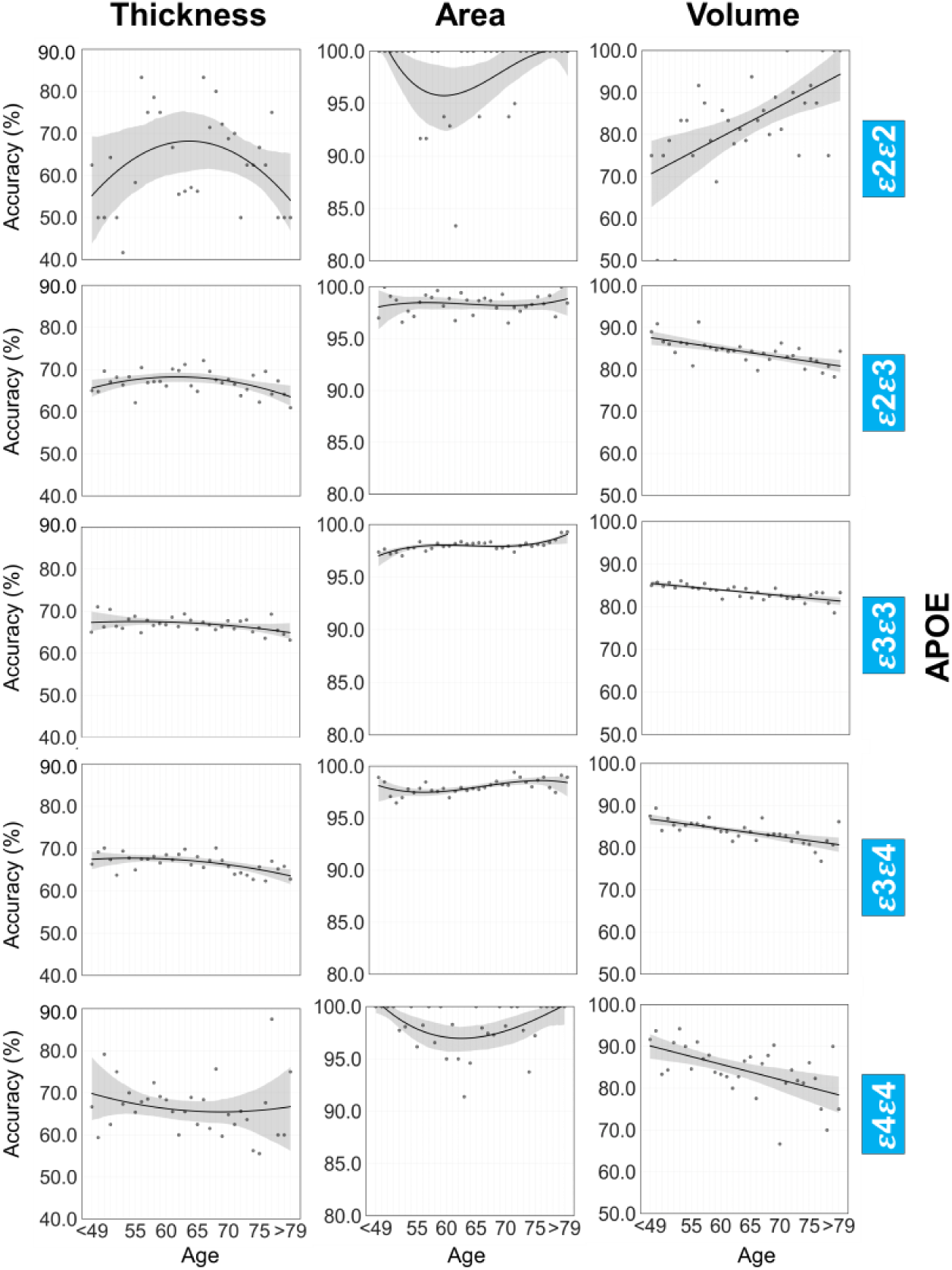
Multivariate brain asymmetry and AD genetic risk factors. APOE ε2 is typically considered to be a protective factor for late-onset AD and APOE ε4 has been considered as a risk factor. Note that, considering the limited sample size of each age year of below 49 years old in the dataset, samples with an age of younger than 49 years old were grouped together when calculating the statistics (e.g., prediction accuracy). Similarly, samples of above 79 years old were also grouped.

**Fig. S6.**
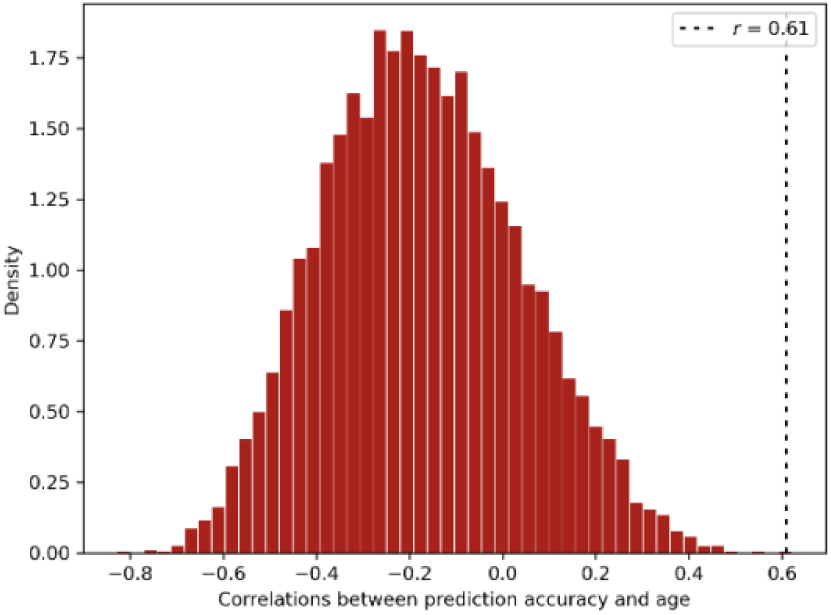
Correlation distribution for random samples with the same sample size as ε2ε2 group (N=194). Random samples were selected from those whose were not ε2ε2 carries. The correlation between prediction accuracy and age was calculated for each time of sampling. This calculation was repeated for 10000 times. The dashed line indicates the observed correlation between age and the prediction accuracy in ε2ε2 group, which was higher than 99.99% correlations of the random samples.

**Fig. S7.**
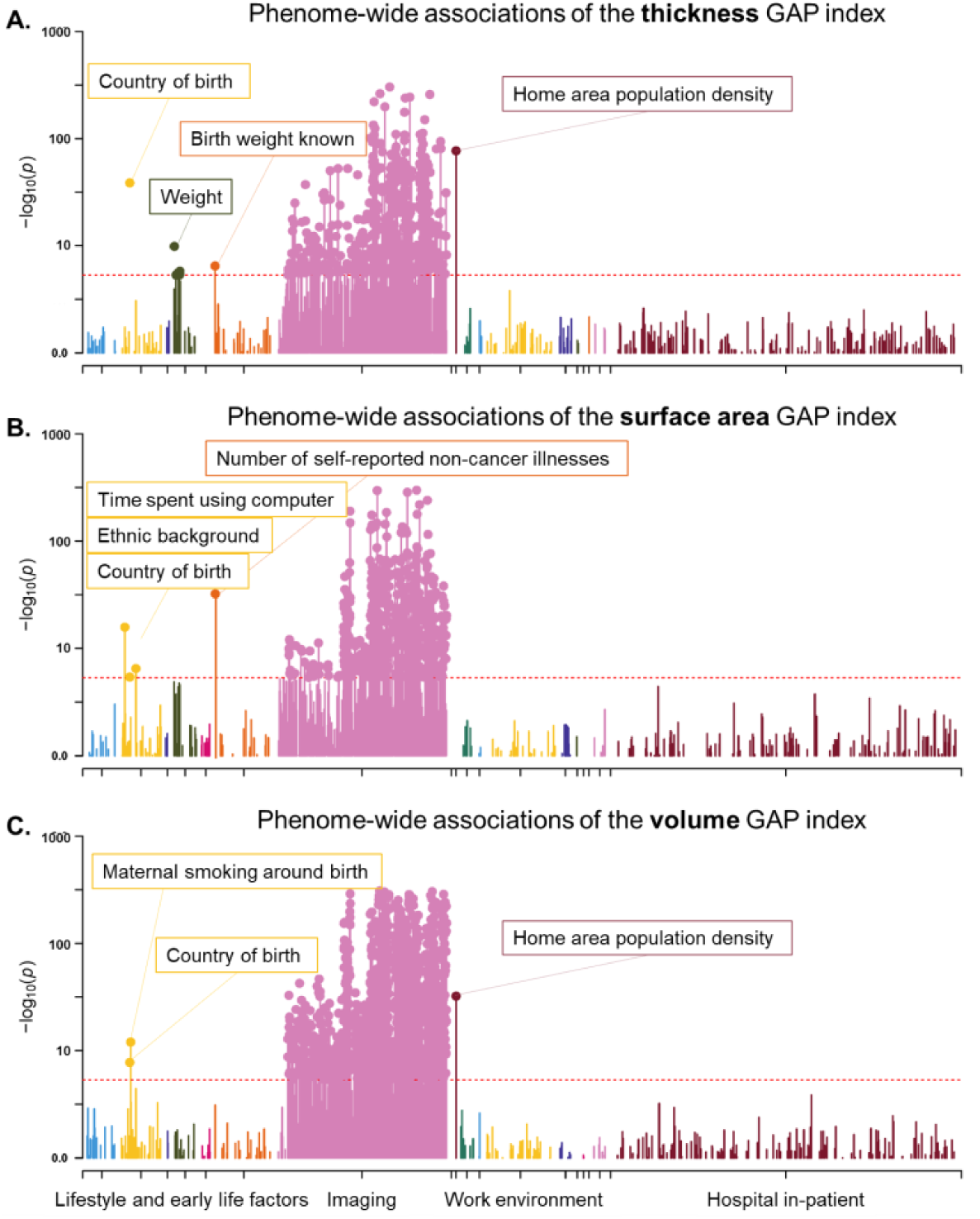
Manhattan plots for the phenome-wide associations of the GAP index from the global brain asymmetry models. Phenome-wide association results for the model of cortical thickness (A), surface area (B), and volume (C). Red lines indicate the Bonferroni corrected threshold of *p* = 4.64e−06. Significant associations are annotated with labels.

**Fig. S8.**
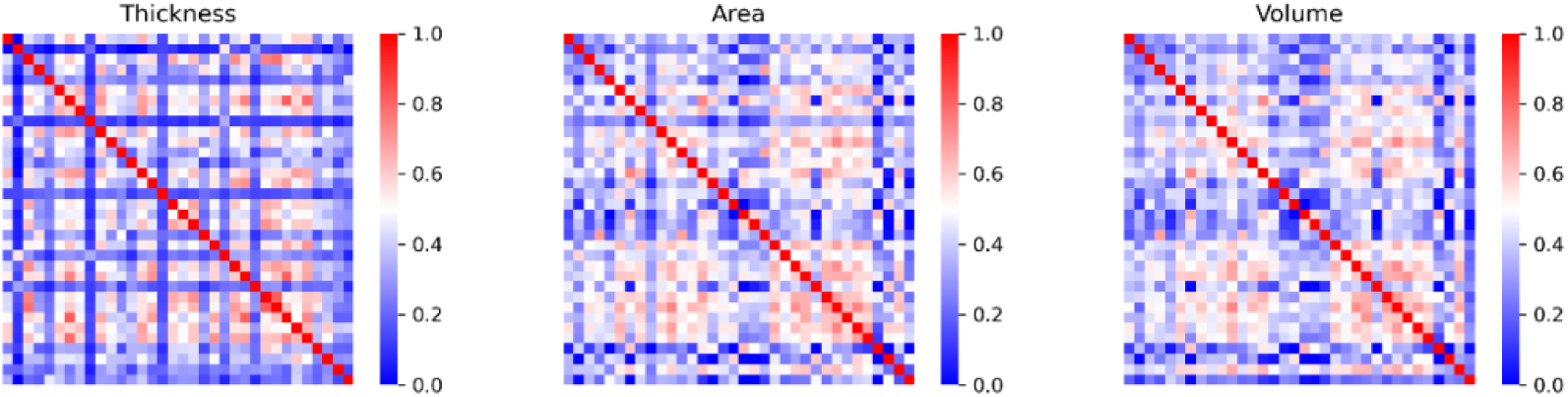
Correlation maps for the regional structural measures in the HCP. Correlations of thickness: mean=0.36, std=0.18, max=0.81; correlations of area: mean=0.40, std=0.17, max=0.75; volume: mean=0.40, std=0.16, std=0.73. No correlations higher than 0.9 (i.e., a rule of thumb for collinearity) were observed for either of the three metrics.

